# Biodiversity of Tropical Forests

**DOI:** 10.1101/088534

**Authors:** A. Tovo, S. Suweis, M. Formentin, M. Favretti, Jayanth R. Banavar, S. Azaele, A. Maritan

**Author notes:** These authors contributed equally to this work (A. Tovo, S. Suweis).

## Abstract

The quantification of tropical tree biodiversity worldwide remains an open and challenging problem. In fact, more than two-fifths of the global tree population can be found either in tropical or sub-tropical forests^1^, but species identities are known only for ≈ 0.000067% of the individuals in all tropical forests^2^. For practical reasons, biodiversity is typically measured or monitored at fine spatial scales. However, important drivers of ecological change tend to act at large scales. Conservation issues, for example, apply to diversity at global, national or regional scales. Extrapolating species richness from the local to the global scale is not straightforward. Indeed, a vast number of different biodiversity estimators have been developed under different statistical sampling frameworks^3–7^, but most of them have been designed for local/regional-scale extrapolations, and they tend to be sensitive to the spatial distribution of trees^8^, sample coverage and sampling methods^9^. Here, we introduce an analytical framework that provides robust and accurate estimates of species richness and abundances in biodiversity-rich ecosystems, as confirmed by tests performed on various *in silico*-generated forests. The new framework quantifies the minimum percentage cover that should be sampled to achieve a given average confidence in the upscaled estimate of biodiversity. Our analysis of 15 empirical forest plots shows that previous methods^10,11^ have systematically overestimated the total number of species and leads to new estimates of hyper-rarity^10^ at the global scale^11^, known as Fisher’s paradox^2^. We show that hyper-rarity is a signature of critical-like behavior^12^ in tropical forests^13–15^, and it provides a buffer against mass extinctions^16^. When biotic factors or environmental conditions change, some of these rare species are more able than others to maintain the ecosystem’s functions, thus underscoring the importance of rare species.

---

Tropical forests have long been recognized as one of the largest pools of biodiversity^1^. Global patterns of empirical abundance distributions show that tropical forests vary in their absolute number of species but display surprising similarities in the distribution of populations across species^8,17,18^. A common statistical tool used to describe the commonness and rarity of species in an ecological community is the relative species abundance (RSA), which is a list of species present within a region along with the number of individuals per species^19^. Typically, the RSA is measured at local scales (e.g., in quadrats or transects, see Figure 1), in which the identities of the majority of the individuals living in the area are known. Of course, the sampled RSA can be fit to any desired functional form at that scale. However, to obtain an up-scaled RSA, as would be measured if it were possible to survey an entire forest, it is necessary to incorporate sampling effects, which strongly bias the actual form of the RSA. Generally, an RSA at a given scale leads to an RSA of a different functional form at a smaller scale, thus hindering analytical treatment^20^. Here, we present a theoretical analytical framework to extrapolate species richness from local to global scales. This framework is based on scaling theory and Nachbin’s theorem^21,22^, thus guaranteeing that a given RSA can be approximated to any degree of precision, at least for populations less than some fixed but otherwise arbitrary value, with a linear combination of negative binomial (NB) distributions (see Supplementary Information, sections 1 and 2). A single NB distribution is given by

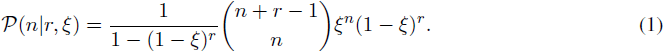

which is normalized so that 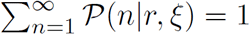, where *r* > 0 and 0 ≤ ξ < 1 are adjustable parameters that control the shape of the RSA. Fisher’s log-series (LS) is obtained as the *r* → 0 limit of equation (1). Owing to sampling effects, a small sample of a forest will have several singleton species with just one individual. In contrast, a larger sample may exhibit an internal mode, and this attribute is well captured by the NB distribution with effective parameters. The NB distribution can be derived from first principles on the basis of biological processes (see Methods). When there are no correlations in the placement of the individuals, the functional form of the NB remains the same after upscaling but with effective parameters, and this remains an accurate approximation even when there are correlations, as validated by in silico experiments. This two-parameter functional form is versatile (see Figure 2 and Supplementary Information, section 1) and is able to adequately fit the RSAs of diverse ecosystems such as tropical forests and coral reefs^23^ and accounts for the effects of density dependence^17,24,25^ on birth rates. The continuum version of the NB, i.e., the gamma distribution, is also the stationary state of a model that captures the temporal turnover of species, an important aspect of tropical tree dynamics^26^. The great advantage of the NB distribution is that it is form-invariant under up-scaling (see Methods). Under up-scaling, a single NB distribution (equation (1)) retains the same value of the parameter *r*, which is scale invariant, and a new effective value of the other parameter, ξ. The same holds true for a linear combination of NB distributions with different values of *r* and the same ξ. Here again, after up-scaling, the original *r* values are retained, and an effective ξ parameter is obtained.

**Figure 1:**
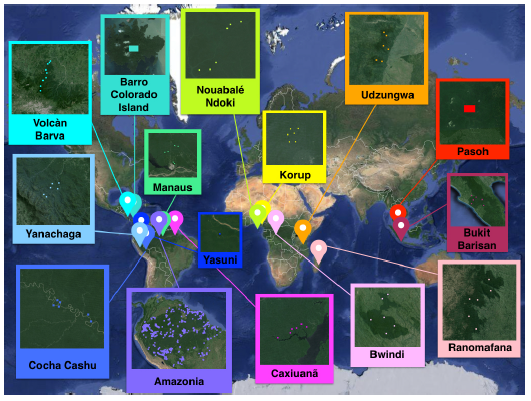
The challenge of estimating global tropical biodiversity. A map depicting the 15 forests of our dataset for which the coordinates of each subplot (squares) are known. Our goal was to deduce the biodiversity of each entire forest on the basis of the very limited knowledge in the marked dots (see Table 2 and Supplementary Information, section 3, for a more detailed description of the dataset).

**Figure 2:**
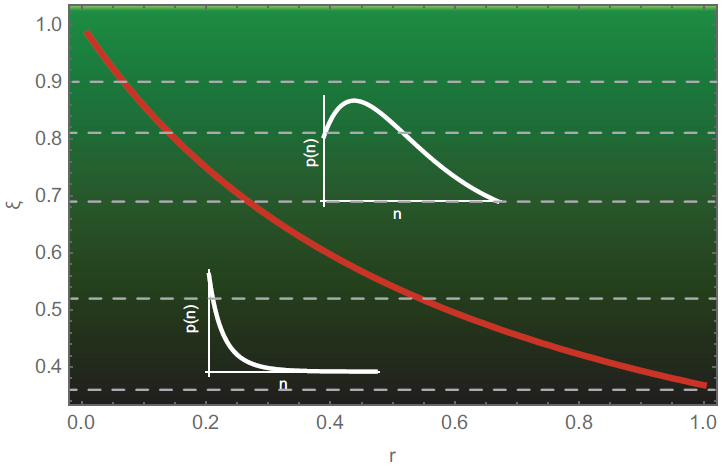
Versatility of the NB distribution. The NB distribution is a two-parameter distribution that shows self-similarity and can display both monotonic log series-like behavior (in the limit *r* → 0 the NB tends to the LS distribution) and a unimodal shape, as a function of the scaling parameter ξ. The red line represents the analytical threshold separating these two cases. The RSA distribution, especially at large scales or with increasing sampling effort^29^, often displays an interior mode that cannot be captured by the LS distribution but can be described by the NB. The NB distribution naturally arises as the steady-state RSA of an ecosystem undergoing generalized dynamics of birth, death, speciation, and migration processes (see Methods). Finally, any discrete probability distribution such as the RSA can be approximated to any degree of accuracy by a suitable linear combination of NBs that retains the self-similarity feature (see Methods). An example is shown of how the parameter ξ of the NB increases as the area of the forest doubles. Starting from ξ=0.36, as the area doubles, the ξ value moves to the value corresponding to the successive (dashed) horizontal line in the upward direction.

We now formulate our analytical framework on the basis of the following two steps. 1) Begin with data on the abundances of *S\** species within a given region covering a fraction *p\** of the whole forest, i.e., 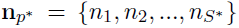 describes the abundances of the sampled species. 2) Use a linear combination of a suitable number of NBs with the same 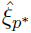 and different values of *r* to fit the empirical RSA at the desired degree of accuracy. This method is guaranteed to be effective according to Nachbin’s theorem^21,22^. The RSA distributions at different scales have the same functional form as the RSA at the scale *p\**, and only the value of the parameter ξ change as a function of the scale (see Methods). Thus, we obtain an analytical solution of the upscaled RSA at scale *p* from the data at scale *p\** in terms of the equation 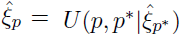, defining 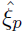 in terms of *p*, *p\** and 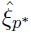. Therefore, using the RSA at scale *p\**, a maximum likelihood method is used to estimate the parameters of the RSA distribution, and the upscaling equations (see Methods) are used to predict the species richness of the entire forest, *p* = 1. The framework resembles the renormalization group technique in critical phenomena in which the behavior of a system at different scales is described in terms of equations for the model parameters, similarly to the proposal here^12^. By using our framework (the NB framework in the following), we were able to generate accurate and robust predictions for computer-generated forests (see Table 1) and for 15 empirical tropical forests (Supplementary Information, section 3)(Figure 1 and Table 2). We also found that the previous method^10,11^ based on the LS distribution (LS method in the following) has several drawbacks (discussed in Supplementary Information, section 4) and leads to a significant overestimation of rare species and consequently of the total forest biodiversity (Tables 1 and 2). We first compared the results of our method applied to a computer-generated forest with species abundances extracted from a log-normal distribution and spatial correlations according to a modified Thomas process (see Supplementary Information, section 5) when fitting the RSA with an LS and with a single or a linear combination of two NBs. In this *in silico* experiment, we fixed the number of species (in this case S = 5000) and their abundance distribution *a priori* and then generated the forest accordingly. We then randomly sampled only a small fraction, *p\**, of individuals and attempted to predict *S* by using only this partial information. The predicted biodiversity is shown in Table 1 within the NB framework for different spatial correlations of forest trees. We found that even though the original forest had a log-normal RSA entangled with spatial correlations, a single NB or a linear combination of two NBs led to surprisingly good predictions and systematically outperformed the LS method; this result was also true for computer-generated forests with different RSAs (see Supplementary Information, section 5). Using more than one NB introduced extra parameters in the fitting and thus made it possible to increase the accuracy of the prediction of the number of species at the global scale. However, even with a single NB, the predictions were quite good, as shown in Table 1. For the forest data, we chose to work with just a single NB that can be derived from basic ecological processes, such as birth, death and migration^8,23^ (see Methods). This simplification was also validated by the results of our *in silico* experiments. This approach permits an exact analytical treatment with just two parameters for the fitting of the sampled RSA and the prediction of the corresponding RSA and of the total number of species at the global scale.

**Table 1:** Estimation of the total biodiversity of a computer-generated forest by using a single negative binomial distribution or a linear combination of two negative binomial distributions. The results of the LS method are also shown for comparison. We generated a forest composed of 5000 different species with a log-normal RSA distribution of mean *μ* = 5 and standard deviation *τ* = 1. The individual trees were located according to a modified Thomas process (see Supplementary Information, section 5) with two distinct clustering coefficients. We then sampled 5% of the global area and applied our method. The prediction of the number of species using the NB framework with just one negative binomial was already quite good (error > 2%). The introduction of two additional fitting parameters, which are necessary when using a linear combination of two negative binomials, improved the estimates (error < 0.2%). In contrast, the LS method overestimated the number of species (error > 56%).

|  | **LS** |  | **one NB** |  | **two NB** |  |
| --- | --- | --- | --- | --- | --- | --- |
|  | High-clustered | Low-clustered | High-clustered | Low-clustered | High-clustered | Low-clustered |
| $S_{pred}$ | 7838 | 9036 | 5095 | 5067 | 4995 | 5011 |

Our results for tropical forests around the globe are presented in Table 2 (and also Supplementary Information, section 6). We found that the LS method^10,11^ systematically overestimated the number of species at the largest scale. Only for the Yanachaga Chimillen National Park were the two estimates with NB and LS essentially the same. The discrepancies in the estimate increased to approximately 10% for Amazonia and Barro Colorado (BCI), reach 30 – 40% for Pasoh and Bukit Barisan and ranged between 72% and 152% for the remaining 10 forests. The errors in our estimates are also given in Table 2. A further prediction of our framework is shown in Table 3 Methods and Supplementary Information, section 7), which indicates, for each forest, the amount of sampling (p_pred_% - second column) necessary to achieve biodiversity predictions with errors below approximately 5% within a 95% confidence interval and its ratio with respect to the actual sampling (third column). Apart from BCI, Caxiuana, Korup, Manaus, Pasoh, Volcan and Yasuni, for which our estimates of the total number of species already had an error below approximately 5%, we predict that all the other forests would require further sampling. Amazonia, for example, would need approximately double the current sampling, Cocha and Nouable approximately tenfold, and Bwindi, Udzungwa and Yanachaga several hundred-fold their current sampling.

**Table 2:** Predicting Biodiversity in Tropical Forests. Predicted total number of species, *S_pred_*, in each of the 15 tropical forests, determined by using the NB framework with a single negative binomial for fitting the sampled RSA, corresponding to a fraction *p\** of the entire forest with *N\** trees, in which only *S\** species are seen. Standard errors were computed by propagating the errors in the fitting parameters of the RSA, obtained by the bootstrapping method, and of *S\**, determined as follows: for each dataset, we created the corresponding predicted forest at the global scale by generating *S_pred_* numbers distributed according to a negative binomial of parameters (*r, ξ*), and we sampled the p*% of the list of individuals, as in the original data.

| Forest | $S^*$ | $N^*$ | $p^*\%$ | $S_{pred}$ (NB) | $S_{pred}$ (LS) |
| --- | --- | --- | --- | --- | --- |
| AMAZONIA | 4962 | 553949 | 0.00016 | $13602 \pm 711$ | 14984 |
| BARRO COLORADO | 301 | 222602 | 3.20513 | $366 \pm 15$ | 419 |
| BUKIT BARISAN | 340 | 14974 | 0.00169 | $471 \pm 40$ | 1020 |
| BWINDI | 128 | 18490 | 0.01813 | $163 \pm 15$ | 288 |
| CAXIUANA | 386 | 32701 | 0.01818 | $437 \pm 14$ | 915 |
| COCHA CASHU | 489 | 16640 | 0.00035 | $731 \pm 63$ | 1674 |
| KORUP | 226 | 17427 | 0.00473 | $282 \pm 23$ | 591 |
| MANAUS | 946 | 38933 | 0.06000 | $1016 \pm 14$ | 2242 |
| NOUABALE NDOKI | 110 | 7196 | 0.00143 | $125 \pm 8$ | 316 |
| PASOH FOREST RESERVE | 927 | 310520 | 0.35714 | $1193 \pm 36$ | 1590 |
| RANOMAFANA | 269 | 34580 | 0.01463 | $336 \pm 22$ | 620 |
| UDZUNGWA | 109 | 18447 | 0.00302 | $146 \pm 20$ | 269 |
| VOLCAN BARVA | 392 | 44439 | 0.02025 | $448 \pm 16$ | 895 |
| YANACHAGA | 209 | 2041 | 0.00372 | $802 \pm 211$ | 802 |
| YASUNI | 481 | 13817 | 0.61100 | $565 \pm 20$ | 974 |

**Table 3:** Sampling targets for forest percentage cover. Using our results on up-scaled forest biodiversity, it was possible to estimate the percentage p_pred_% of the forest that must be sampled to achieve an estimation error of approximately 5%. We derived these values by creating the predicted forest at the global scale (we generated *S_pred_* numbers according to a negative binomial of parameters *r* and ξ) and sampled it at increasingly larger scales until the desired accuracy in the estimation of the global biodiversity was reached (see Supplementary Information, section 7 for more details). The last column indicates how much extra sampling is needed (if the number is greater than 1) to reach 5% precision.

| Forest | $p_{pred}\%$ | $p_{pred}/p^*$ |
| --- | --- | --- |
| AMAZONIA | 0.0003 | 1.875 |
| BARRO COLORADO | 3 | 1 |
| BUKIT BARISAN | 0.05 | 18 |
| BWINDI | 5 | 386 |
| CAXIUANA | 0.01 | 0.55 |
| COCHA CASHU | 0.003 | 8.57 |
| KORUP | 0.02 | 1.06 |
| MANAUS | 0.02 | 0.17 |
| NOUABALE NDOKI | 0.015 | 10.5 |
| PASOH FOREST RESERVE | 0.5 | 1.4 |
| RANOMAFANA | 0.1 | 6.84 |
| UDZUNGWA | 1.5 | 497 |
| VOLCAN BARVA | 0.02 | 0.25 |
| YANACHAGA | 1 | 269 |
| YASUNI | 0.3 | 0.49 |

We estimated the number of hyper-rare species, defined as species with fewer than 1000 individuals, and hyper-dominant species, defined as the most abundant species contributing approximately 50% of the total population^10^. The numbers of hyper-rare and of hyper-dominant species (see Table 4) have also been previously^10,11^ overestimated and underestimated, respectively. In fact, the asymptotic value of Fisher’s *α* is strongly biased when a very small fraction of the forest is sampled (typically < 1%) (see Supplementary Information, section 4). Moreover, we found that the hyper-rarity phenomenon is an emergent pattern in tropical forests, which may characterize biodiversity hotspots^27^. Our framework provides a possible explanation for this phenomenon, considering the observed hyper-rarity as a manifestation of criticality in tropical forests^12,13^. Indeed, the parameters of the NB distributions that provided the best predictions of the up-scaled biodiversity in tropical forests all clustered around 0 < *r* < 0.7 and ξ ≈ 1. This result was somewhat surprising, because there are neither theoretical nor biological reasons that might explain why tropical forests in different geographical locations and with differing species richness should have abundances distributed across species in a very similar manner. However, closer examination of the form of the NB distribution revealed that the relative fluctuation of abundances, i.e., 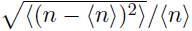 diverged as ξ → 1 and *r* → 0 (see Figure 3 and Supplementary Information, section 8). Thus, parameter values close to this region allow an ecosystem to have the highest heterogeneity in its abundance distribution. The points shown in Figure 3 correspond to the parameter values obtained for the 15 forests. A physical system such as water and vapor, in the vicinity of its critical point, is characterized by density fluctuations that become very large, with droplets of water and bubbles of gas of all sizes thoroughly interspersed, and the system appears the same at different scales, i.e., it is self-similar^12^. This scale invariance confers on the system an exquisite sensitivity to certain types of external perturbations or disturbances whose effects are realized at long distances. The observed abundance fluctuations suggest that tropical forests may be critical systems and may be relatively reactive to disturbances^14,15^ and able to adapt optimally to new external conditions/constraints. Under a given set of environmental conditions, only a few species are best at exploiting the limited resources^28^. Because of environmental fluctuations, these conditions may not continue to be advantageous for the existing very few abundant species. However, a large pool of species serves as a reservoir of new opportunities and responses and as a buffer against newly changed conditions^28^. According to this view, hyper-rarity is essential for an ecosystem to maintain its functions and react promptly to changes: rare species provide the key to an ecosystem’s future^16^.

**Figure 3:**
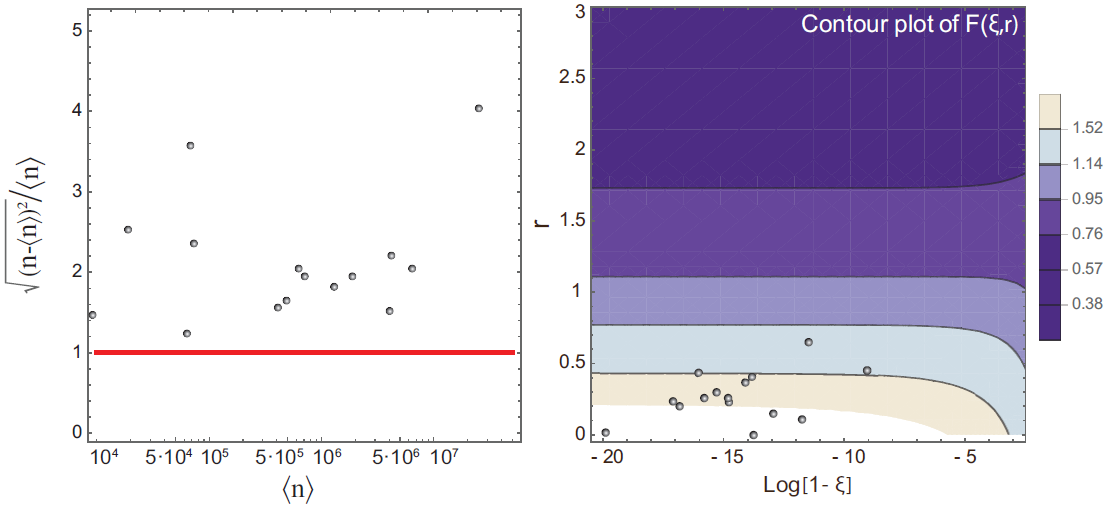
Tropical forests are poised in the vicinity of criticality. A) Plot of the relative fluctuations of species abundances, 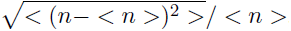, in linear scale versus abundances < n > at the logarithmic scale. The black points denote the predicted values for each of the 15 tropical forests listed in Table 2 at the global scale, and the red curve is the line of equation *y* = 1. All values are located above this line, thus indicating that the relative fluctuations in abundance are considerable for all the forests. B) Contour plot of the relative fluctuation of abundances for a negative binomial RSA 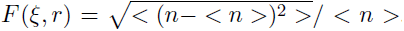. The black points represent the pair (*r*, log[1 - ξ]), where *r* and ξ are the predicted parameters for each forest of our dataset after up-scaling at the global scale. These points are all located in the region of the parameter space around which the function *F(ξ, r)* diverges, i.e., ξ ≈ 1 and 0 < *r* < 0.7.

**Table 4:**
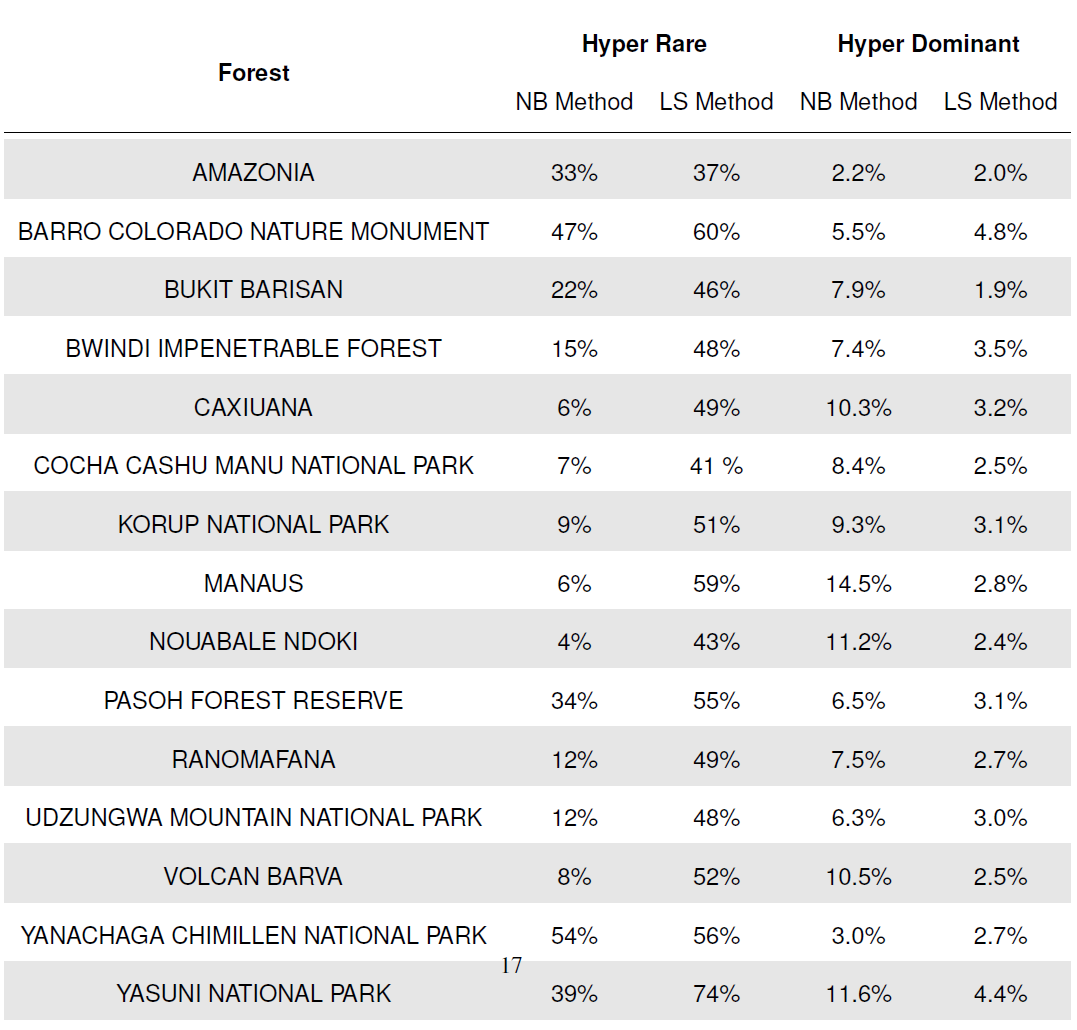
Fisher’s paradox^2^. Hyper-rare species (defined as species with fewer than 1000 individuals^10,11^) and hyper-dominant species (the most abundant species, accounting for ≈ 50% of the total number of individuals) percentages were predicted in the whole area of each tropical forest obtained by applying both the NB and LS methods. Interestingly, we found that by using our NB method, the number of hyper-rare species in most of the forests was drastically reduced with respect to the LS method, thus suggesting that the extremely high value of hyper-rare species predicted in previous studies^10,11^ is an artifact of the LS method. Nevertheless, we found that the hyper-rarity phenomenon is a genuine emergent pattern in tropical forests.

## Methods

### 1 Upscaling negative binomials

Here, we chose the negative binomial (NB) distribution in equation (1) as the RSA. Apart from its simplicity and versatility, we chose this form for our analysis for four reasons:

1. Any discrete probability distribution such as the RSA can be approximated to any degree of accuracy by a suitable linear combination of NBs (see section 2 of the Supplementary Information for a proof and discussion). We made the parsimonious choice of a single NB function because it suffices to approximately describe the available tropical forest data, as discussed in the main text.
2. The NB distribution arises naturally as the steady-state RSA of an ecosystem with sufficiently weak inter-species interactions and undergoing generalized dynamics of birth, death, speciation, and immigration to and emigration from the surrounding metacommunity (see next section).
3. In the limit of *r* → 0, the NB becomes the well-known Fisher log-series (LS), which has been widely used to describe the patterns of abundance in ecological communities. Of course, because of the flexibility of choosing *r* to be non-zero, the NB distribution is always more versatile than the LS. The RSA distribution, especially at large scales or with increasing sampling effort^29^, often displays an interior mode that cannot be captured by an LS distribution. Indeed, the Fisher log-series is not sufficiently flexible^20^ to capture different patterns of RSAs ^8,23,30–34^, especially a count of the rare species in tropical forests (see Supplementary Information, section 4). To assess whether the increased reliability of the NB method with respect to the LS method is due only to the introduction of the additional parameter *r*, we used the Akaike information criterion, which shows that the NB is the preferred model for all tropical forests of our dataset except one for which *r* is very close to zero.
4. Finally, if one chooses two contiguous patches with NB as RSA distributions characterized by the same parameters *r* and ξ ≡ ξ_1/2_ and combines the two, remarkably, the resulting larger patch is also characterized by an NB distribution with the same scale-invariant value of *r* and a new scale-dependent parameter, ξ, given by the analytical expression in equation (3) below with *p* = 1/2. This special form-invariant property of the NB distribution, albeit with a scale-dependent parameter, makes it particularly well suited for our extrapolation studies.

Denoting the fraction of the sampled area of a forest by *p*, one finds (see Supplementary information, section 1) that the total number of species at the largest scale (*p* = 1) is related to the number of species at scale *p*, *S_p_*, by

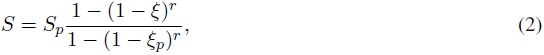

where ξ_*p*_ and *r* are the fitted parameters of the NB of the RSA distribution at scale *p*. As noted above, *r* is scale invariant and hence independent of *p*, whereas the parameter ξ at the largest scale, *p* = 1, is given by

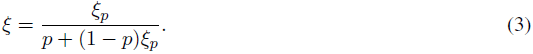

We demonstrate that the framework holds exactly when species are spatially uncorrelated (see Supplementary Information, section 9). However, our in *silico* experiments indicated that the framework is robust even in the presence of spatial correlations and for different sampling methods (Supplementary Information, section 5). Finally, we tested the lack of importance of spatial correlations in predicting singletons in tropical forest data by generating null models in which the species labels of trees are shuffled (Supplementary Information, section 9).

## 2 Stochastic model leading to negative binomial RSA

As previously stated, the NB distribution arises naturally as the steady-state RSA of an ecosystem that undergoes simple birth and death dynamics under the neutral hypothesis, wherein species are not distinguishable, and all inter-specific interactions are neglected^23^.

Indeed, let 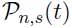 be the probability that, at time *t*, species *s* have exactly *n* individuals, where *s* ∈ {1,…,*S*}. We assume that the population dynamics of each species is governed by two terms, *b_n,s_* and *d_n,s_*, which are the birth and death rates for species *s* with *n* individuals. The master equation regulating the evolution of 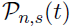 for *n* ≥ 0 is then

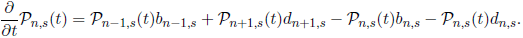

The above equation is also valid for *n* = 0 and *n* = 1 if we set 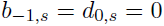. The steady-state solution is

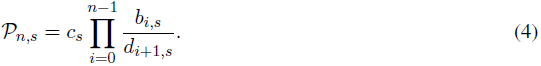

The term *c_s_* is a normalization factor found by imposing 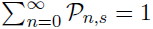.

Let us assume that the birth term in the above equation depends on a density-independent term *b_s_*, which is the per-capita birth rate, and on the term *Y_s_*, which may take into account immigration events or intra-specific interactions:

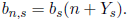

Analogously, let us suppose that the death term depends on a density-independent term *d_s_*, which is the per-capita death rate:

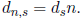

These suppositions are reasonable in ecology. By substituting in eq. (4), we obtain

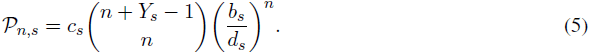

The normalization constant can be easily found by imposing

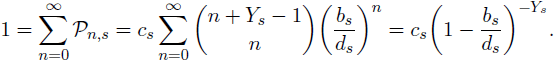

Therefore, the probability that the *s^th^* species has *n* individuals at equilibrium is given by a negative binomial of parameters 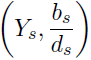:

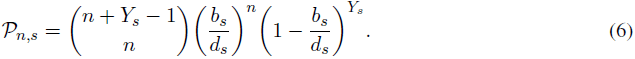

Under the neutral hypothesis, in which all species are considered to be equivalent, we can remove the species index *s* from the above equation, thus obtaining a negative binomially distributed RSA for the ecosystem under study.

## 3 Data availability

We use a global-scale compilation of 1248 local sites collected over 15 forests around the planet on different tropical field stations of the equatorial zone. All data are publicly available. See Supplementary Information for web links and a more detailed description of the dataset.

## 4 Code availability

For numerical simulations performed in this study we used standard commands and programming tools in R/Mathematica. All codes are available upon request.

## Acknowledgements

We are indebted to Steve Hubbell for his insightful comments. The BCI forest dynamics research project was founded by S.P. Hubbell and R.B. Foster and is now managed by R. Condit, S. Lao, and R. Perez under the Center for Tropical Forest Science and the Smithsonian Tropical Research in Panama. Numerous organizations have provided funding, principally the U.S. National Science Foundation, and hundreds of field workers have contributed. All data in this publication were provided by the Tropical Ecology Assessment and Monitoring (TEAM) Network, a collaboration between Conservation International, the Missouri Botanical Garden, the Smithsonian Institution, and the Wildlife Conservation Society, and partially funded by these institutions, the Gordon and Betty Moore Foundation, and other donors.

## Author Contributions

A.T. and S.S. carried out the numerical simulation, the data analysis and performed the figures. A.T, M.F., M.F & A.M. carried out analytical calculations. All the authors contributed to other aspects of the paper and the writing of the manuscript.

## Author Information

Reprints and permissions information is available at www.nature.com/reprints. The authors declare no competing financial interests. Readers are welcome to comment on the online version of the paper. Correspondence and requests for materials should be addressed to S.S. or to A.M.

## References

[1] Crowther, T. et al. Mapping tree density at a global scale. Nature 525, 201–205 (2015).

[2] Hubbell, S. P. Estimating the global number of tropical tree species, and fisher’s paradox. Proceedings of the National Academy of Sciences 112, 7343–7344 (2015).

[3] Bunge, J. & Fitzpatrick, M. Estimating the number of species: a review. Journal of the American Statistical Association 88, 364–373 (1993).

[4] Brose, U., Martinez, N. D. & Williams, R. J. Estimating species richness: sensitivity to sample coverage and insensitivity to spatial patterns. Ecology 84, 2364–2377 (2003).

[5] Mao, C. X. & Colwell, R. K. Estimation of species richness: mixture models, the role of rare species, and inferential challenges. Ecology 86, 1143–1153 (2005).

[6] Wang, J.-P. Z. & Lindsay, B. G. A penalized nonparametric maximum likelihood approach to species richness estimation. Journal of the American Statistical Association 100, 942–959 (2005).

[7] Bunge, J. et al. Estimating population diversity with catchall. Bioinformatics 28, 1045–1047 (2012).

[8] Azaele, S. et al. Statistical mechanics of ecological systems: Neutral theory and beyond. Rev. Mod. Phys. 88, 035003 (2016). URL http://link.aps.org/doi/10.1103/RevModPhys.88.035003.

[9] Chao, A., Colwell, R. K., Lin, C.-W. & Gotelli, N. J. Sufficient sampling for asymptotic minimum species richness estimators. Ecology 90, 1125–1133 (2009).

[10] Ter Steege, H. et al. Hyperdominance in the amazonian tree flora. Science 342, 1243092 (2013).

[11] Slik, J. F. et al. An estimate of the number of tropical tree species. Proceedings of the National Academy of Sciences 112, 7472–7477 (2015).

[12] Stanley, H. E. Scaling, universality, and renormalization: Three pillars of modern critical phenomena. Reviews of modern physics 71, S358 (1999).

[13] Zillio, T., Banavar, J. R., Green, J. L., Harte, J. & Maritan, A. Incipient criticality in ecological communities. Proceedings of the National Academy of Sciences of the United States of America 105, 18714–7 (2008). URL http://www.pubmedcentral.nih.gov/articlerender.fcgi?artid=2596237&too=14]

[14] Hidalgo, J. et al. Information-based fitness and the emergence of criticality in living systems. Proceedings of the National Academy of Sciences 111, 10095–10100 (2014).

[15] Hidalgo, J., Grilli, J., Suweis, S., Maritan, A. & Munoz, M. A. Cooperation, competition and the emergence of criticality in communities of adaptive systems. Journal of Statistical Mechanics: Theory and Experiment 2016, 033203 (2016).

[16] Hull, P. M., Darroch, S. A. & Erwin, D. H. Rarity in mass extinctions and the future of ecosystems. Nature 528, 345–351 (2015).

[17] Volkov, I., Banavar, J., He, F., Hubbell, S. & Maritan, A. Density dependence explains tree species abundance and diversity in tropical forests. Nature 438, 658–61 (2005). URL http://www.ncbi.nlm.nih.gov/pubmed/16319890.

[18] McGill, B. J. et al. Species abundance distributions: moving beyond single prediction theories to integration within an ecological framework. Ecology letters 10, 995–1015 (2007).

[19] MacArthur, R. On the relative abundance of species. American Naturalist 25–36 (1960).

[20] Azaele, S. et al. Towards a unified descriptive theory for spatial ecology: predicting biodiversity patterns across spatial scales. Methods in Ecology and Evolution 6, 324–332 (2015).

[21] Nachbin, L. Sur les algebres denses de fonctions differentiables sur une variete. Comptes rendus hebdomadaires des seances de l'Academie des sciences 228, 1549–1551 (1949).

[22] Llavona, J. G. Approximation of continuously differentiable functions. North-Holland mathematics studies 130, vii–x (1986).

[23] Volkov, I., Banavar, J. R., Hubbell, S. P. & Maritan, A. Patterns of relative species abundance in rainforests and coral reefs. Nature 450, 45–49 (2007). URL http://dx.doi.org/10.1038/nature06197.

[24] Janzen, D. H. Herbivores and the number of tree species in tropical forests. American naturalist 501–528 (1970).

[25] Clark, D. A. & Clark, D. B. Spacing dynamics of a tropical rain forest tree: evaluation of the janzen-connell model. American Naturalist 769–788 (1984).

[26] Azaele, S., Pigolotti, S., Banavar, J. R. & Maritan, A. Dynamical evolution of ecosystems. Nature 444, 926–928 (2006). URL http://dx.doi.org/10.1038/nature05320.

[27] Myers, N., Mittermeier, R. A., Mittermeier, C. G., Da Fonseca, G. A. & Kent, J. Biodiversity hotspots for conservation priorities. Nature 403, 853–858 (2000).

[28] Grilli, J., Suweis, S. & Maritan, A. Growth or reproduction: emergence of an evolutionary optimal strategy. Journal of Statistical Mechanics: Theory and Experiment 2013, P10020 (2013).

[29] Chisholm, R. A. Sampling species abundance distributions: resolving the veil-line debate. Journal of theoretical biology 247, 600–607 (2007).

[30] Chave, J. Neutral theory and community ecology. Ecology Letters 7, 241–253 (2004). URL http://doi.wiley.com/10.1111/j.1461-0248.2003.00566.x.

[31] Magurran, A. Species abundance distributions: pattern or process? Functional Ecology 19, 177–181 (2005).

[32] Chave, J., Alonso, D. & Etienne, R. S. Comparing models of species abundance. Nature 441, E1 (2006).

[33] Magurran, A. E. Ecological diversity and its measurement (Springer Science & Business Media, 2013).

[34] Matthews, T. J. & Whittaker, R. J. Neutral theory and the species abundance distribution: recent developments and prospects for unifying niche and neutral perspectives. Ecology and evolution 4, 2263–2277 (2014).

